# Automated CAR-T Cell Manufacturing: from Research to Point of Care production

**DOI:** 10.64898/2026.09.08.748871

**Authors:** Caroline Boudousquié, Laeticia Dutoit, Fabien Jammes, Luca Bellosta, Emilie Nallet, Yann Pierson, Jonathan H. Esensten, Nicola Vannini, Maria Morbidelli, Valérie Widmer, Martin Pedard, Melita Irving, Jimmy Maillard, Romain Vuillefroy de Silly, Denis Migliorini

## Abstract

**Background:** Point-of-care manufacturing of CAR-T cells could reduce cost and improve patient access, but few closed, automated systems support both flexible process development and GMP production.

**Methods and results:** We describe LimCORE, a single-use consumable for cell incubation and processing, implemented in LimGROW (manual system) and LimONE (automated system). In LimGROW, T cells expanded 246 +/-18-fold over 10 days at optimized seeding density, versus 89 +/-9-fold in a G-Rex control, with similar viability (>90%) and phenotype. Process parameters directly transferred to LimONE, where automated, closed buoyancy-based CD3+ selection achieved 90.3 +/-3.5% purity and 59.2 +/-17.0% recovery in under two hours. Using this workflow, we manufactured CD19 CAR-T cells in an automated 7-day LimONE process requiring 155 minutes of operator intervention. LimONE manufactured CAR-T cells reached 42.6 +/-2.4-fold expansion (532 +/-29 × 10⁶ cells at harvest), with 43.6 +/-6.0% transduction efficiency and viability similar to G-Rex controls. CD4+/CD8+ ratios, CD8+ differentiation subsets, and cytotoxicity were comparable between platforms, while LimONE-manufactured cells showed increased metabolic fitness. Across CAR-T manufacturing runs on the LimONE, no hardware or software failures occurred, and automated liquid transfers and volume concentration stayed within ±5% accuracy.

**Conclusions:** These data show that LimCORE supports T cell expansion, selection, and complete CD19 CAR-T manufacturing with performance similar to existing platforms while reducing operator time, across R&D and GMP scale.

## Introduction

Point-of-care manufacturing of effector T cell products has the potential to decrease cost and improve patient access, even in patients who may have access to commercial CAR T therapies [1,2]. However, there are few commercially available point-of-care manufacturing systems that can be easily and effectively deployed by hospital or regional health systems. The CliniMACS Prodigy® is a market leader in point-of-care CAR T manufacturing [3–7] and has been successfully deployed across the world for both academic and commercial CAR-T manufacturing using single-use tubing sets and standardized manufacturing protocols. The Cocoon® Platform is a more recently introduced system that uses a single-use cartridge for CAR-T manufacturing [8]. Both platforms are controlled by a proprietary software that requires some degree of specialized technical knowledge and/or manufacturer support to modify the protocol for a given process, though the Cocoon® software is marketed as offering more direct, on-site protocol editing than the CliniMACS Prodigy®. Neither platform offers a dedicated R&D tool, so process development must be carried out directly on the full manufacturing system, driving up the cost of process development and making tech transfer from R&D to GMP manufacturing cumbersome. A variety of stand-alone bioreactors with limited additional cell processing functionality are also commonly used in point-of-care manufacturing, including the G-Rex^®^, Sefia™, and Xuri™ bioreactors [2,9].

There are widespread efforts for many hospital and health systems to establish their own manufacturing capacity. In one survey, of FACT/JACIE accredited institutions, 40/45 had the intention to do in-house CAR-T manufacturing [10]. Despite this enthusiasm for local point-of-care manufacturing, significant barriers remain, including limited access to classified cleanroom space, lack of access to viral vectors, and elevated regulatory expectations for clinical trials involving CAR-T with well-established targets, and commercial CAR options [11].

Here we demonstrate the functions and performance of LimCORE, a single-use consumable with a proprietary geometry that enables cell incubation, centrifugation, and precise volume control within the same closed unit. LimCORE is implemented in two tools: LimGROW, an R&D scale rotation platform, and LimONE, a closed and automated system designed for GMP manufacturing. Rotating the LimCORE supports cell incubation on both LimGROW and LimONE, while LimONE additionally exploits its geometry to enable centrifugation-based cell processing with embedded CO_2_ and temperature control. This allows LimONE to perform buoyancy-based cell selection, incubation, transduction, media addition and exchange, and washing entirely within a single unit, without opening the system or transferring cells between vessels. We demonstrate that LimCORE achieves performance in line with established conventional closed-system culture devices across these processing and incubation steps, combining flexibility to adapt to diverse process workflows with ease of use and reduced operator time. We further show end-to-end manufacturing of a CD19 CAR T cell product on LimONE, including cell selection, transduction, expansion, and harvest.

## Materials and methods

### PBMC isolation

Peripheral blood mononuclear cells (PBMCs) were obtained from healthy donor buffy coats. Buffy coats are anonymized, commercial byproducts of routine voluntary blood donation, purchased from an accredited blood transfusion center (Transfusion Interrégionale CRS, Epalinges, Switzerland). Donor informed consent for research use of surplus material is obtained by the center at the time of donation. As the material is fully de-identified and commercially sourced, no additional institutional ethics committee approval was required for this study. Buffy coats were processed fresh upon receipt, and PBMCs were isolated by Ficoll density-gradient centrifugation using Ficoll-Paque (Cytiva, USA; Cat. 17144003) and SepMate PBMC tubes (STEMCELL Technologies, Canada; Cat. 85460).

### CAR construct and lentiviral production and titration

Lentiviruses were produced in 293T cells (ATCC) by polyethylenimine (PEI Max, Polysciences)-mediated transfection at a 3:1 PEI-to-plasmid mass ratio. The plasmid mixture comprised, in a 3:2:1 ratio, the transfer lentivector encoding an anti-CD19 second-generation CAR with a CD8α hinge, 4-1BB co-stimulatory and CD3ζ signalling domains (CAR-T-2-L332-Bz, Creative Biolabs), the packaging plasmid pCMVR8.74 (Addgene #22036; RRID:Addgene_22036; gift from D. Trono) and the envelope plasmid pMD2.G (Addgene #12259; RRID:Addgene_12259; gift from D. Trono). Supernatants were harvested two days post-transfection, concentrated by ultracentrifugation (24,000g for 2 h), and stored at –80 °C. As previously described with modification [12], lentiviral content was titrated based on CD19 CAR positivity by flow cytometry after serial dilution on Jurkat E6-1 cells (ATCC) in the presence of 10 µg/mL protamine sulfate (Sigma-Aldrich, P4020).

### PBMC seeding, activation and expansion

After isolation or thawing, PBMCs were seeded at 1–2 × 10⁶ cells/mL in 2–10 mL working volumes. Activation was performed with TransACT (1:200, Miltenyi Biotec, Germany; Cat. 130-111-160) in T-VIVO medium (Lonza, Basel, Switzerland; Cat. BP12-970Q) supplemented with 300 IU/mL Interleukin-2 (IL-2, AcroBiosystems, Delaware, USA; Cat. GMP-L02H14). Cultures were maintained at 37 °C and 5% CO₂, with media replenishment as needed. The same culture conditions were run in G-Rex 6M flasks (Wilson and Wolf, USA, Cat.# 80660M) with 5 × 10⁶ cells seeded according to the manufacturer’s recommendation.

### Manual T cell selection

T cells were isolated from fresh PBMCs by negative selection using the Human T Cell Isolation Kit (Akadeum Life Sciences, Ann Arbor, MI, USA; Cat. 13210-120) according to the manufacturer’s instructions. Briefly, PBMCs were resuspended in complete cell culture medium at 2–4 × 10⁸ cells/mL in a sterile transfer bag. Air equivalent to 20% of the bag volume was added aseptically to facilitate mixing. Non-T cells were labelled by aseptic addition of the Human T Cell Leukopak Biotin Antibody Cocktail targeting CD14, CD16, CD19, CD20, CD36, CD56, CD123 and CD235ab (1 mL per 1 × 10⁹ cells) followed by 10 min on an orbital shaker at room temperature. Streptavidin-conjugated BACS™ microbubbles (12.5 mL per 1 × 10⁹ cells) were then added aseptically following thorough resuspension to achieve a homogeneous suspension, and samples were mixed by orbital shaking for 15 min at room temperature. Additional complete medium was added as needed, and the bag was suspended from a hanger for 10 min to allow buoyant microbubble-bound non-T cells to rise to the surface. Untouched T cells were recovered by gravity drainage of the negatively-selected cellular fraction into a sterile collection vessel.

### Automated T cell selection in LimONE

Human T cells were isolated from PBMCs by negative selection using the Human T Cell Leukopak Isolation Kit (Akadeum Life Sciences; Cat. 13210-221) or the Human T Cell Isolation Kit (Cat. 13210-120), integrated into an automated closed-system workflow on the LimONE platform (Limula SA, Lausanne, Switzerland). The full process was performed under aseptic conditions with steps guided by the LimONE software. Following tubing-set installation and priming with PBS (Corning, Glendale, AZ, USA; Cat. 21-040-CV) and 0.5% human serum albumin (Gemini Bio-Products, Cat. GEMB800-121-050), PBMCs were transferred aseptically in a sterile bag under a biological safety cabinet (BSC) and welded to the tubing kit prior to loading. The biotinylated antibody cocktail (CD14, CD16, CD19, CD20, CD36, CD56, CD123, CD235ab) was injected aseptically via syringe through a sterile connector and mixed with PBMCs for 15 min at room temperature. The antibody-labelled cell suspension was then transferred to a microbubbles (MB) bag prepared under BSC and welded to the tubing kit. Streptavidin-conjugated BACS™ microbubbles were added following thorough resuspension, and the PBMC/antibody/microbubble mixture was incubated with bag shaking for 15 min at room temperature. Gravity-based sedimentation was performed by hanging the bag on the LimONE hook for 20 min, allowing buoyant microbubble-bound non-T cells to rise to the surface. Untouched T cells were then automatically retrieved in LimONE and sampled for cell counting via syringe. Isolated T cells were subsequently seeded while the excess T cell fraction was collected into a sterile bag.

### T cell seeding and activation

Enriched T cells were seeded at 0.5 × 10⁶ cells/mL. Activation was performed using TransACT at 1:200 (Miltenyi Biotec, Germany, Cat. 130-111-160) in T-VIVO medium (Lonza; Cat. BP12-970Q) supplemented with 300 IU/mL IL-2 (Akron, Boca Raton, FL, USA; Cat. AR1045-0010). IL-2 supplementation of the culture medium was automated using the LimONE system. IL-2 (1 × 10⁶ IU in 10 mL; Akron, USA, Cat. AR1045-0010) and the culture medium bag were connected to the LimONE disposable kit. Briefly, 100 mL of culture medium was transferred into the rotor, followed by 3 mL of IL-2 solution (300’000 IU). The mixture was homogenized within the rotor and returned to the culture medium bag of 1 L, resulting in a final concentration of 300 IU/mL IL-2. The supplemented medium was used throughout the 7-day process with sealing/welding and storage at 2–8 °C when not actively used.

### T cell transduction in LimONE

One day after seeding 12.5 × 10⁶ T cells, transduction was performed by adding hCD19 CAR-T lentivirus (UNIL, Lausanne, Switzerland) at a multiplicity of infection (MOI) of 1 in combination with LentiBoost 1:400 (Revvity, USA; Cat. SB-A-LF-901-02) before returning to static expansion until day 3. The virus was added to 1 mL of T-VIVO medium and transferred into a 5 mL syringe. The syringe was then connected to the LimONE using a MicroCNX sterile connector (CPC Biotech, USA; Cat. CNX170LFHT) and the volume transferred into the rotor.

### T cell expansion in LimONE

On day 3, a media addition was performed with twice the seeding volume of T-VIVO medium supplemented with 300 IU/mL IL-2. The bag of T-VIVO and IL-2 was prepared on day 0 for the whole process; when not used it was stored at 2–8 °C and warmed at room temperature before addition. Dynamic incubation at 14 rpm, 37 °C, 5% CO₂ was then started. On day 5, a media addition with T-VIVO and 300 IU/mL IL-2 was performed to reach a cell concentration of 1 × 10⁶/mL (if the maximum volume of the rotor was not reached). If maximum incubation volume was reached (300 mL), an in-situ centrifugation was performed to remove 80% of the supernatant. Cells were then resuspended in the maximum volume of medium and dynamic incubation resumed at 14 rpm, 37 °C, 5% CO₂. On day 7, the cell suspension was harvested by transferring all the rotor contents into an empty 600 mL bag and cells were counted. On days 3 and 5, sampling for cell counting was performed via the LimONE syringe kit: the rotor was placed in an upright position and a 1 mL sample was taken in a 5 mL syringe, the tubing was welded, and the sample was transferred into a conical tube for counting.

### T cell activation, transduction and expansion in the G-Rex control

On day 0, 5 × 10⁶ cells were added to one well of a G-Rex 6M plate (Wilson and Wolf, USA, Cat.# 80660M) containing T-VIVO medium with 300 IU/mL IL-2 and activated with TransACT (1:200). On day 1, transduction was performed with the hCD19 CAR-T lentiviral vector (UNIL) at MOI 1 in combination with LentiBOOST^TM^ at 1:400 (Revvity, USA). On day 3, the well was topped up with T-VIVO and 300 IU/mL IL-2. On day 5, a cytokine refresh with 300 IU/mL IL-2 was performed on the total volume. On day 7, the supernatant was removed and the cell suspension was harvested and counted.

### CAR-T cell cytotoxicity assay

Cytotoxic activity was assessed using live-cell imaging on the Incucyte S3 system (Sartorius). GFP-expressing NALM6 human B cell leukemia cells (ATCC; Cat. CRL-3273) were used as targets and seeded at 2 × 10⁴ cells/well in 96-well flat-bottom plates. CAR-T cells and non-transduced T cells were added at effector-to-target (E:T) ratios of 3:1(normalized to CAR-T^+^ cells). Co-cultures were performed in complete T cell medium at 37 °C with 5% CO₂ for 72 h. GFP fluorescence confluence was acquired every 2 h. Cytotoxicity was calculated as 100 × (1 – raw GFP confluence at time t / mean GFP confluence of target-only control wells at time t).

### SeaHorse assay

Oxygen consumption rates (OCRs) were measured using a Seahorse XFe96 Bioanalyzer (Agilent). Cells were resuspended in Seahorse XF Base Medium supplemented with 10 mM glucose, 1 mM sodium pyruvate, and 2 mM glutamine (pH 7.4) and maintained at 37 °C. They were seeded at 2 × 10^5^ cells per well in Cell-Tak-coated (22.4μg/mL) Seahorse XFe96 microplates. Injection ports were loaded with 1μM oligomycin, 2μM carbonyl cyanide-p-trifluoromethoxyphenylhydrazone (FCCP), and 0.5μM rotenone/antimycin A. During sensor calibration, plates were incubated for 45 min at 37 °C in a non-CO₂ incubator. OCR data were analyzed using Seahorse Wave software (version 2.4.3). Spare respiratory capacity (SRC) was calculated from a Seahorse Mito Stress Test as the difference between the maximal OCR after FCCP treatment and the basal OCR. T cells were plated at 2 × 10⁵ cells per well on Cell-Tak-coated Seahorse XFe96 microplates, as previously described [13], and OCR values are reported per well at this fixed seeding density. Each donor was measured in three technical replicate wells. Each donor was measured in three technical replicate wells (n = 4 donors, except LimONE Untransduced, n = 2 donors). Data are presented as mean ± SEM of donor-level values (n = 4 donors, except LimONE Untransduced, n = 2).

### Antibodies and flow cytometry

Cell phenotyping was performed at multiple stages of the manufacturing process. For pre- and post-T cell selection products (day 0), the panel included CD3-FITC (clone OKT3, BioLegend; Cat. 317305), CD4-PerCP-Vio700 (M-T466, Miltenyi; Cat. 130-113-819), CD8α-BV421 (RPA-T8, BioLegend; Cat. 301036), CD19-SB600 (HIB19, ThermoFisher; Cat. 63-0199-42), CD14-AF594 (HCD14, BioLegend; Cat. 325630), CD56-APC (5.1H11, BioLegend; Cat. 362503), CCR7-PE/Cy7 (G043H7, BioLegend; Cat. 353226) and CD45RA-BV510 (HI100, BioLegend; Cat. 304142). For harvested T cell products (day 7), the panel comprised CD19 CAR-T Biotin (Miltenyi; Cat. 130-129-550) revealed with anti-Biotin-PE (REA746, Miltenyi; Cat. 130-111-068), CD3-FITC, CD4-PerCP-Vio700, CD8α-BV421, CCR7-PE/Cy7 and CD45RA-BV510.

Cells were washed and resuspended in flow cytometry (FC) buffer (PBS supplemented with 2% FBS [Clearline, North Wales, PA, USA; Cat. 509503] and 2 mM EDTA [VWR; Cat. E177-100]). Dead cells were excluded with Zombie NIR fixable viability dye (BioLegend; Cat. 423106). Fc receptors were blocked with Human TruStain FcX (BioLegend; Cat. 422302) combined with the viability dye for 15 min at 4 °C. The antibody cocktail was added and incubated for a further 20 min at 4 °C. Cells were fixed with BioLegend fixation buffer (Cat. 420801) for 10 min at room temperature, washed in FC buffer, and stored until acquisition on a CytoFLEX S or LX flow cytometer (Beckman Coulter). Data were analysed using CytExpert v2.6 (Beckman Coulter).

The following fluorochrome-conjugated antibodies were used for tetramethylrhodamine methyl ester (TMRM) analysis: CD3–BV711 (clone UCHT1, BioLegend; Cat. 300464), CD4–BV605 (clone OKT4, BioLegend; Cat. 317438), CD8–APC (clone SK1, BioLegend; Cat. 344722), CCR7–BV421 (clone G043H7, BioLegend; Cat. 353208), CD45RO-PE-Cy7 (clone UCHL1, BioLegend; Cat. 304230), CD45RA-Spark-UV387 (cloneHI100, BioLegend; Cat. 304180), CD62L–PerCP-Cy5.5 (clone DREG-56, BioLegend; Cat. 304824).

For TMRM staining (MedChemExpress, CAS No.: 115532-50-8), cells were incubated with 10 nM of TMRM for 15 min at 37°C, 5% CO_2_. Samples were read at the Cytek® Aurora and analyzed using FlowJo^TM^10.

### Automated cell counting

Cell counting and viability were determined using the LUNA-FX7 automated cell counter (Logos Biosystems, Anyang, South Korea) with AO/PI staining (Logos Biosystems; Cat. F23001) and dual-fluorescence read-out.

### Accuracy of automated liquid transfers

To evaluate the accuracy of volumetric transfers performed in the LimONE, two target volumes were assessed: 50 mL and 100 mL. For each condition, transfers were performed using water at a flow rate matching the target volume (50 mL/min for 50 mL transfers, and 100 mL/min for 100 mL transfers). Each volume was transferred three times from bag to LimCORE and three times from LimCORE to bag, for a total of six replicates per condition.

The mass of the receiving bag was automatically recorded by the integrated LimONE software via an onboard scale, with density assumed to be 1 g/mL (i.e., 1 mg = 1 µL). The measured volume was then compared to the target volume, and the relative error was calculated as follows:

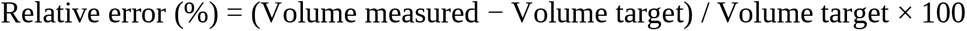

A transfer was considered accurate if the relative error remained within ±5% of the target volume.

### Accuracy of centrifugation-based volume concentration

The accuracy of the end-point volume reached by the LimONE centrifugation process was evaluated using two different starting volumes (350 mL and 700 mL) with a fixed target concentrated volume of 50 mL. Six and four centrifugation runs were performed for the 350 mL and 700 mL conditions, respectively. At the end of each run, the remaining volume in the rotor was measured manually using a pipette and compared to the target volume. The relative error was calculated as described above.

### Graphical rendering and statistical analysis

Data are presented as mean, with each symbol representing an independent donor unless otherwise stated in the figure legends. Comparisons between two independent groups were performed using an unpaired t-test with Welch’s correction. Comparisons across more than two groups were performed using ordinary one-way ANOVA, and comparisons across two independent variables using ordinary two-way ANOVA; both were followed by Tukey’s multiple comparisons test. Statistical analyses were performed using GraphPad Prism (version [11.1]). Statistical significance is indicated as *p < 0.05, **p < 0.01, ***p < 0.001, ****p < 0.0001; ns, not significant.

## Results

### PBMC expansion in LimGROW

To assess the T cell expansion performance of the LimCORE, a proprietary single-use culture vessel, initial experiments were conducted using the LimGROW system, a simple device in which the LimCORE and its accessories are housed. The LimGROW allows for both static and dynamic incubation of the LimCORE using a single direction of rotation. LimGROW can be used within standard cell culture incubators and biosafety cabinets (Figure 1A). To assess T cell expansion in the LimCORE, 2×10⁶ cells/mL PBMC were seeded, activated with TransACT and expanded in the presence of IL-2. Two different seeding volumes were used to assess lower end volume range (2 mL or 10 mL) and the culture was kept static for 3 days before adding fresh complete media and starting dynamic culture (constant rotation of 14 rpm). Cells were kept in culture for up to 13 days with media addition or exchange (depending on volume) at day 5, 7 and 10. A G-Rex 6M control was run in parallel, with 5×10⁶ PBMC seeded into 5 mL on day 0 and top up of complete media at day 3. At day 13, fold expansion in LimGROW seeded at 10mL was of 132.6 +/- 30.1 (mean +/- SD, n=7) compared to 98.9 +/-17.4 (mean +/- SD, n=5) in the G-Rex control (Figure 1B). Viability was also comparable between LimGROW and G-Rex (90.3 +/-5.4 vs 85.3 +/- 6.8 respectively, mean +/- SD) (Figure 1B). In order to optimize LimCORE performance, the seeding cell concentration was decreased to 1×10⁶ /mL in 10 mL total volume. In those conditions, LimCORE supported greater T cells expansion as we observed a mean of 246.4 +/-18.4 fold expansion (mean +/-SD, n=3) in 10 days. In comparison, the G-Rex control reached 89.3 +/-9.3 fold expansion (mean +/-SD, n=5) in the same time-frame and using same starting material (Figure 1C). Cell phenotype was analyzed at harvest to assess both CD4/CD8 ratio as well as CD8 differentiation based CD45RA and CCR7 expression (Figure 1D). No significant difference between cells cultured in LimGROW compared to the control was observed. The flow cytometry analysis approach is shown supplemental in Figure 1. Taken together, these results show that LimCORE supports successful expansion of T cells with performance comparable to or better than the G-Rex control.

**Figure 1.**
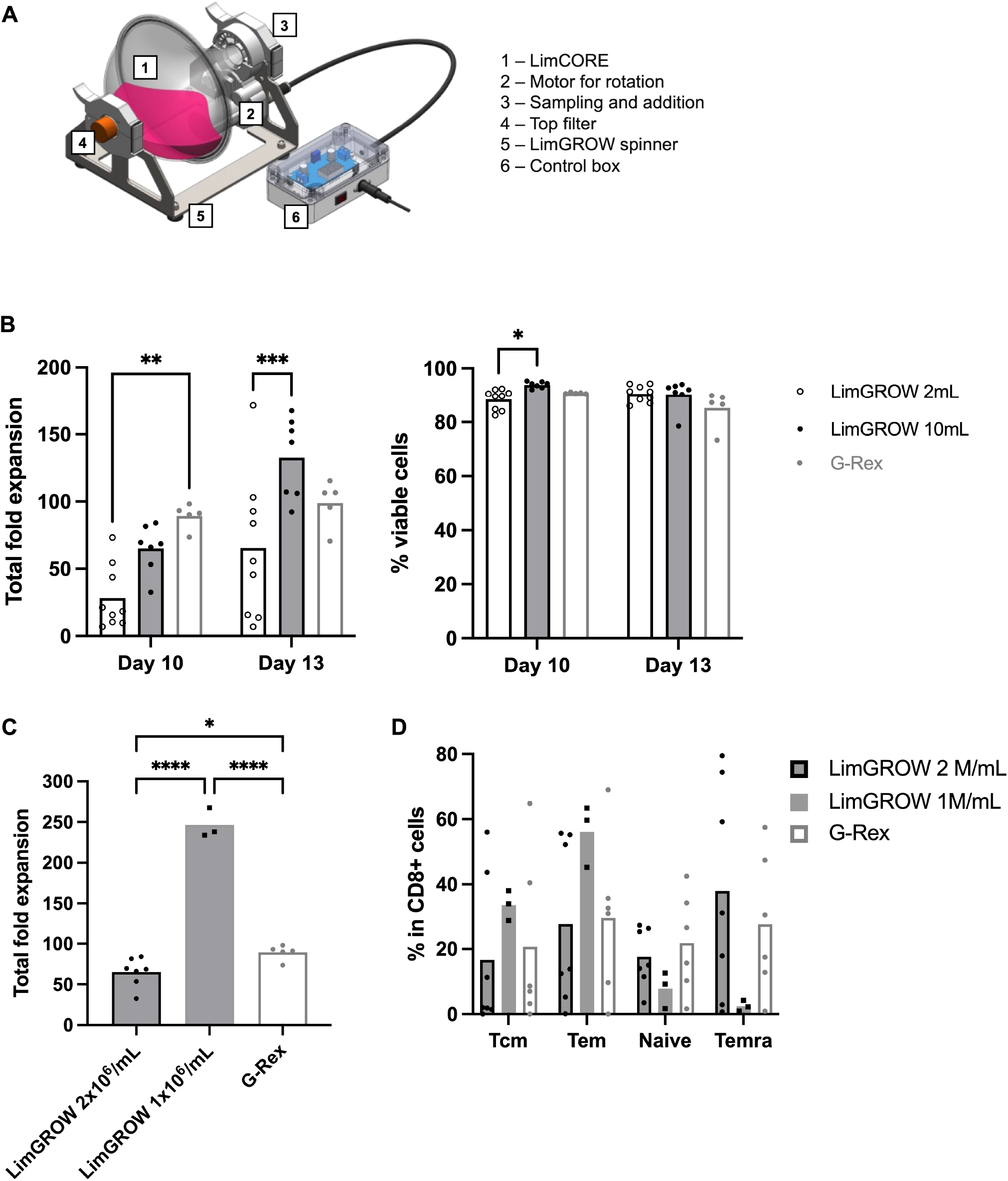
T cells expansion in LimGROW. PBMCs from up to 9 healthy donors were activated and T cells expanded for up to 13 days in LimGROW or G-Rex 6M. (A) LimGROW schematic. (B) LimGROW seeded with PBMCs at 2×10⁶ cells/mL in 2 mL or 10 mL; G-Rex 6M seeded with 5×10⁶ PBMCs. Viability, cell number and fold expansion assessed at day 10 and/or 13. (C) Fold expansion at day 10 in LimGROW (10 mL, 1 or 2×10⁶ PBMCs/mL) vs. G-Rex 6M (5×10⁶ PBMCs). (D) and CD8⁺ memory phenotype (Tcm, Tem, Tnaive, Temra) at harvest. Each symbol represents an independent donor. Statistical significance was determined Two-way (B, D) or one-way (C) ANOVA with Tukey’s multiple comparisons test. *p < 0.05, **p < 0.01, ***p < 0.001, ****p < 0.0001. Not significant difference are not displayed.

### Process transfer from LimGROW to LimONE

LimONE operates using a closed tubing set that includes LimCORE and adds temperature and CO_2_ control, sampling/addition ports, centrifugation for cell processing (e.g., washing or media exchange), and automated liquid handling (Figure 2A). The LimGROW optimized process was transferred to LimONE by using its dedicated protocol authoring software to build an automated protocol from the process parameters defined on LimGROW. After seeding and activating 100×10⁶ of PBMC at day 0 (seeding volume of 100mL), LimCORE remained static from days 0 to 3. At day 3, a media addition was performed and continuous rotation started. 80% media exchange was performed at day 5 and harvest at day 7. At harvest, LimONE reached a higher total expansion than LimGROW (27.2 +/-2.5 fold vs 18.1 +/-2.4 fold, mean+/-SD, n=3) but with a similar cell growth kinetic (Figure 2B). As a control, total fold expansion in the LimONE did not significantly differ from the G-Rex (41.6 +/-17.7 fold, mean+/-SD, n= 14) (Figure 2B). CD4+/CD8+ percentages and T cell subset distributions were similar between the LimGROW and LimONE, consistent with G-Rex control (Figure 2C).

**Figure 2.**
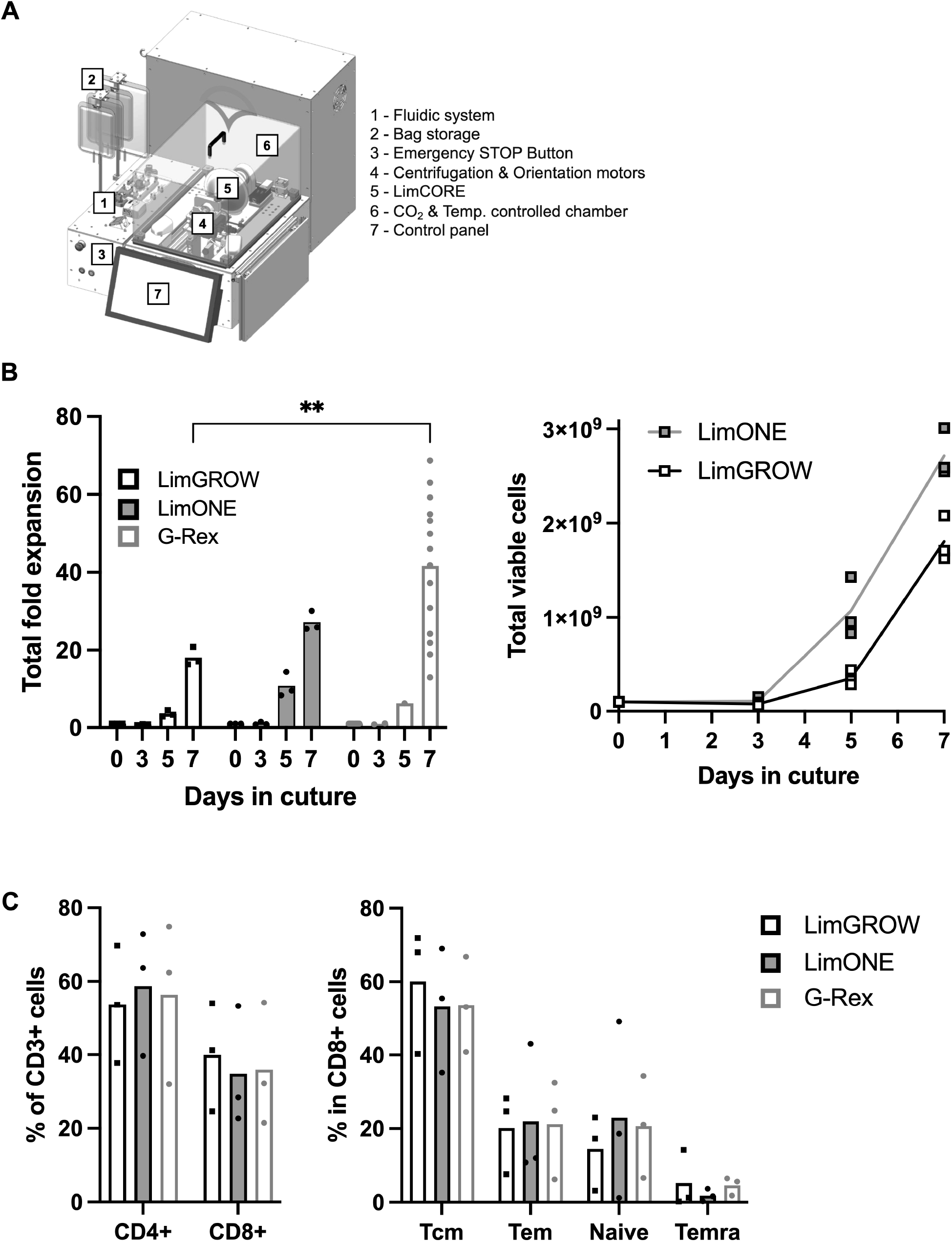
Process transfer to LimONE preserves T cell growth and phenotype. PBMCs from 3 to 14 healthy donors were activated and T cells were expanded for 7 days in LimGROW, LimONE (each seeded with 100×10⁶ cells in a 100 mL seeding volume), or G-Rex 6M (5×10⁶ cells). (A) LimONE schematic. (B) Total fold expansion and total viable cell number are determined by automated AO/PI cell count at day 0, 3, 5, and 7. (C) Phenotype at harvest: CD4⁺ and CD8⁺ proportion of CD3⁺ cells, and CD8⁺ memory subsets (Tcm, Tem, naive, Temra). Each symbol represents an independent donor. Statistical significance was determined using Two-way ANOVA with Tukey’s multiple comparisons test. *p < 0.05, **p < 0.01, ***p < 0.001, ****p < 0.0001. Not significant difference are not displayed.

### Closed, automated buoyancy selection of CD3⁺ T cells in LimONE

The LimONE can also integrate and automate buoyancy-based cell isolation: the isolation itself is performed by commercial reagents (here, Akadeum’s BACS™ microbubbles) [14], while the LimONE handles the process in a closed, automated workflow. This method relies on negative selection: PBMCs are first labelled with a cocktail of biotinylated antibodies that bind non-target lineages (for T-cell isolation: CD14, CD16, CD19, CD20, CD36, CD56, CD123, CD235ab). Streptavidin-conjugated BACS™ microbubbles are then added, binding the biotin-tagged cells. Being buoyant, the microbubbles float the labelled cells to the top of the bag, while the untouched, unlabelled target cells remain in suspension and are recovered as the enriched, negatively-selected fraction. Using a negative selection kit from Akadeum, T cell selection was successfully performed using both a manual (in-bag) and an automated protocol on the LimONE (Figure 3A). The automated workflow was established and tested across eight runs. Following tubing set installation and rotor priming, PBMCs and the biotinylated antibody cocktail were sequentially loaded and mixed into the LimCORE. The labelled cells were then transferred to a bag welded to the tubing set and containing microbubbles before being mixed with streptavidin-conjugated BACS™ microbubbles. Finally, cells and microbubbles are allowed to sediment directly within the bag, floating the labelled non-T cells to the surface. Untouched T cells were transferred back to the LimCORE for sampling and cell counting. The entire selection process was completed in under two hours. Across the eight runs performed, this yielded T cells with 90.3 +/-3.5% CD3 purity (mean +/- SD, n=8) (Figure 3B) and >95% viability (data not shown), with a T cells recovery of 59.2 +/-17.0% (mean +/- SD, n=8) (Figure 3C). Impurities post-selection were routinely under 5%, including very low percentages of CD14+ monocytes in the T cells fraction (Figure 3D).

**Figure 3.**
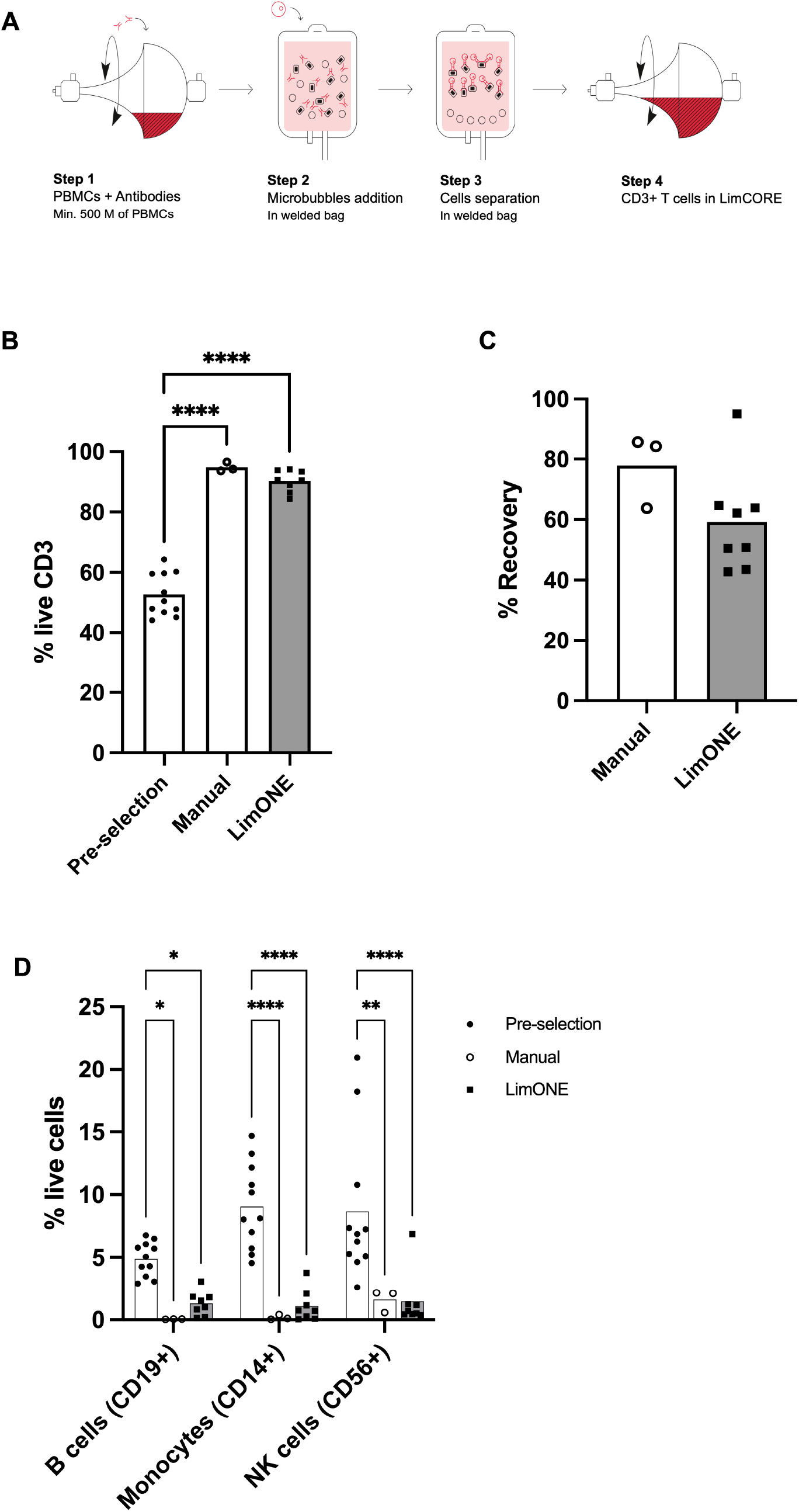
Automated buoyancy based negative CD3⁺ T cells selection. T cell selection was performed on fresh PBMCs from healthy donors using the Akadeum T cell isolation kit with an automated LimONE process (n = 8 donors) or a manual process (n = 3 donors) as per the supplier’s recommendation. (A) Schematic of the four-step automated selection process. (B) Percentage of CD3⁺ T cells before and after selection, determined by flow cytometry. (C) T cell recovery, calculated based on %CD3⁺ T cells and cell count. (D) Impurities characterized by flow cytometry (B cells, CD19⁺; monocytes, CD14⁺; NK cells, CD56⁺) across conditions. Each symbol represents an independent donor. Statistics: (B) ordinary one-way ANOVA and (D) ordinary two-way ANOVA, both with Tukey’s multiple comparisons test; (C) unpaired t-test with Welch’s correction. *p < 0.05, **p < 0.01, ****p < 0.0001; non-significant differences are not displayed.

### From selection to harvest: automated CAR-T manufacturing in LimONE

To demonstrate automated cell and gene therapy manufacturing, we developed a 7-day CD19 CAR-T protocol spanning automated T cell isolation, activation, transduction, culture, and harvest. Operator intervention time was assessed across the complete 7-day CAR-T manufacturing process, from set-up through harvest (Table 1). Summed across all steps, total operator intervention time was approximately 155 minutes (approximately 2.6 hours). By comparison, the CliniMACS Prodigy® has been reported to require approximately 6 hours of hands-on operator time for a comparable manufacturing process [15]. LimONE’s total operator intervention time across the full 7-day process is therefore estimated to be less than half of Miltenyi’s CliniMACS Prodigy® reported hands-on time.

**Table 1.**
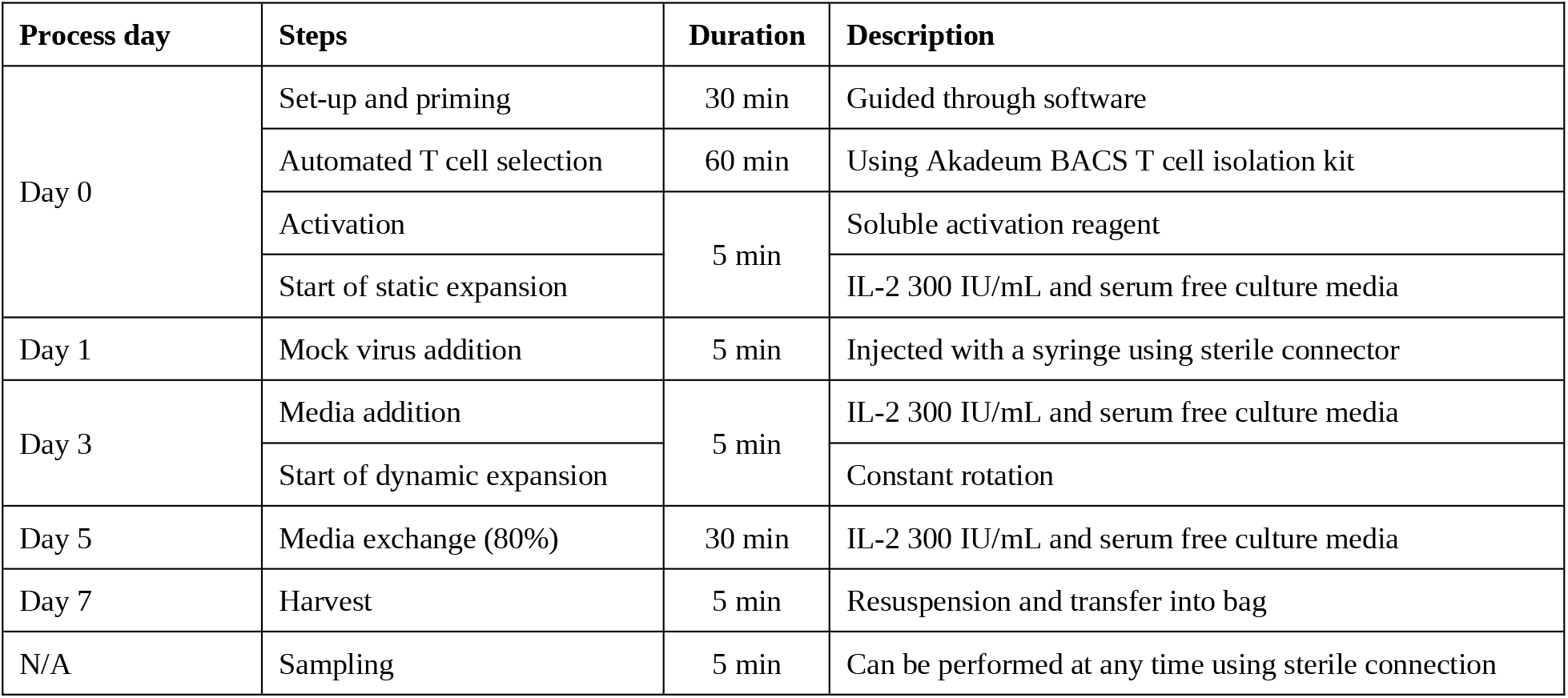
LimONE CAR-T manufacturing process timeline. Process steps, hands-on duration, and description for each day of the 7-day manufacturing process. Sampling can be performed at any time point using sterile connection.

| Process day | Steps | Duration | Description |
| --- | --- | --- | --- |
| Day 0 | Set-up and priming | 30 min | Guided through software |
|  | Automated T cell selection | 60 min | Using Akadeum BACS T cell isolation kit |
|  | Activation | 5 min | Soluble activation reagent |
|  | Start of static expansion |  | IL-2 300 IU/mL and serum free culture media |
| Day 1 | Mock virus addition | 5 min | Injected with a syringe using sterile connector |
| Day 3 | Media addition | 5 min | IL-2 300 IU/mL and serum free culture media |
|  | Start of dynamic expansion |  | Constant rotation |
| Day 5 | Media exchange (80%) | 30 min | IL-2 300 IU/mL and serum free culture media |
| Day 7 | Harvest | 5 min | Resuspension and transfer into bag |
| N/A | Sampling | 5 min | Can be performed at any time using sterile connection |

Complete CAR-T manufacturing runs were performed on the LimONE using material from four healthy donors. We used a CD19 CAR lentiviral vector at a low multiplicity of infection (MOI = 1), with a mock-transduced control run in parallel on a separate LimONE for each donor. Total fold expansion (Figure 4A) and viable cell numbers (Figure 4B) were compared at harvest between transduced and untransduced T cells. With a total input cell number on day 0 of 12.5 × 10⁶ cells in 25 mL of medium, total cell numbers on day 7 in transduced LimONE runs reached 532 +/-29 × 10⁶ cells (mean +/- SD, n=4) (Figure 4B). Total fold expansion in the LimONE (42.6 +/-2.4-fold, mean +/- SD, n=4) was not significantly different from the G-Rex control (49.2 +/-9.9-fold, mean +/- SD, n=4) (Figure 4A). Transduced cells showed a trend toward lower fold expansion (42.6 +/-2.4 fold for LimONE vs 49.2 +/-9.9 fold for G-Rex, mean +/- SD, n=4) compared to untransduced cells (51.8 +/-7.0 fold for LimONE vs 59.1 ±8.3 fold for G-Rex, mean +/- SD, n=4) on both platforms. No differences in cell viability were observed at day 7 (Figure 4C) but transduction efficiency showed a trend toward being higher in the LimONE (43.6 +/-6.0%, mean +/- SD, n=4) compared to the G-Rex control (35.8 +/-6.1%, mean +/- SD, n=4), although this difference was not statistically significant (Figure 4D).

**Figure 4.**
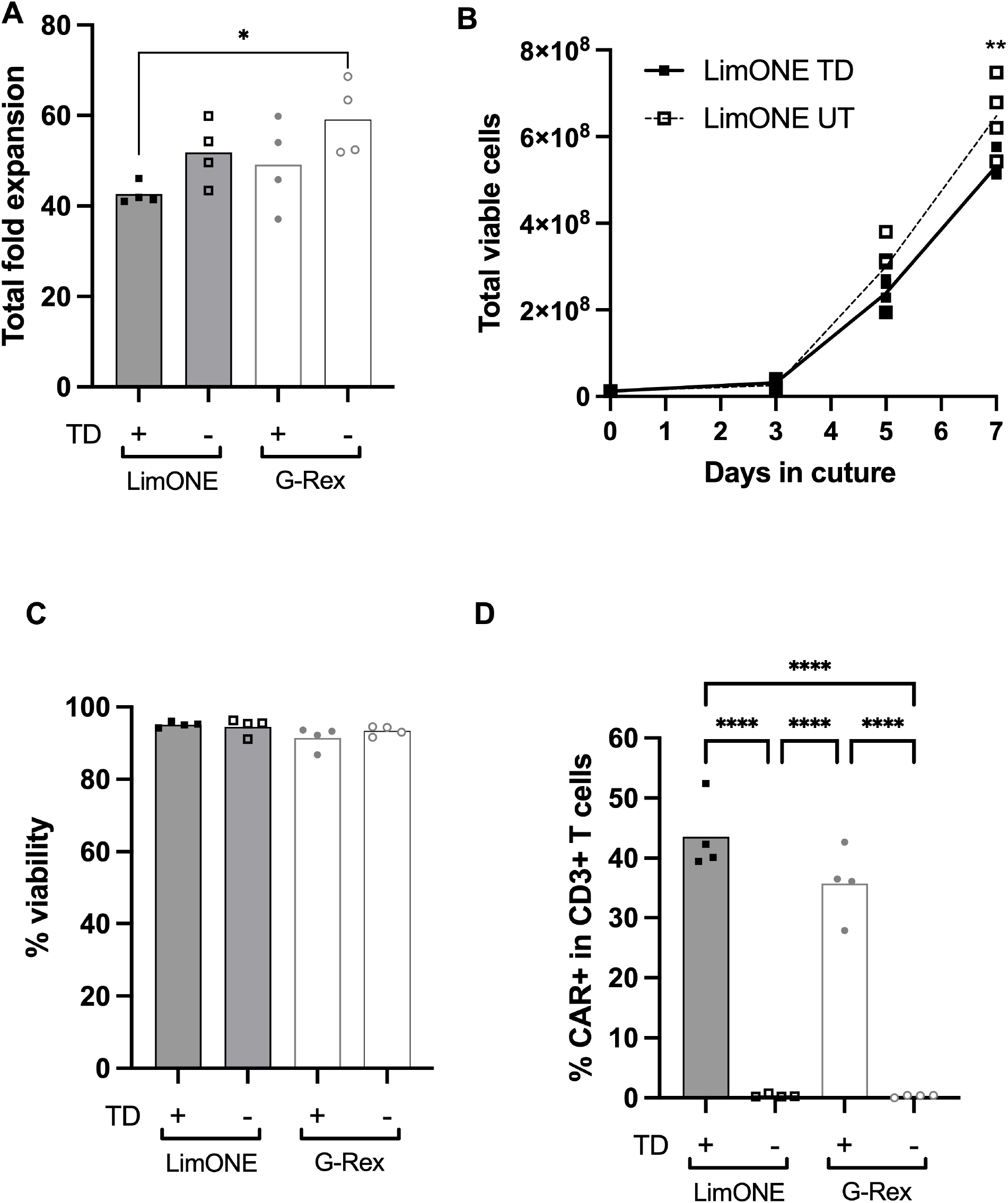
7 days CAR-T process performance in LimONE. Selected T cells were seeded in LimONE at 0.5×10⁶ cells per mL in 25mL before being activated at day 0 and transduced (TD+) or mock-treated (TD−) at day 1 using a CD19 CAR lentiviral vector at a MOI of 1 before being expanded until day 7. As a control, G-Rex were seeded with 5×10⁶ T cells before activation (day 0), transduction (day 1) and expansion. (A) Total fold expansion was determined at day 7 based on cell count. (B) Total viable cell number and (C) cell viability were determined using automated cell count and AO/PI dyes. (D) CAR positivity was determined by flow cytometry (% CD19 CAR⁺ within CD3⁺) at harvest (day 7). Each symbol represents an independent donor (n = 4). Statistics: (A, C, D) one-way ANOVA and (B) two-way ANOVA all with Tukey’s multiple comparisons test. *p < 0.05, **p < 0.01, ****p < 0.0001.

### Characterization of CAR-T cells manufactured in the LimONE

End-of-process phenotyping showed comparable CD8⁺ T cell differentiation (Figure 5A) as well as comparable CD3, CD4 and CD8 percentages (Supplementary Figure 2B) between the LimONE and the G-Rex control. LimONE-manufactured CAR-T cells exhibited potent, sustained killing activity over a 72-hour co-culture time course, comparable to the G-Rex control (Figure 5B). To determine the metabolic fitness of engineered T cells, we measured mitochondrial membrane potential (TMRM mean fluorescence intensity), spare respiratory capacity (SRC), and real-time oxygen consumption rate (OCR) in cells expanded in G-Rex or LimONE, with or without lentiviral transduction. Mitochondrial membrane potential, assessed by TMRM MFI, was comparable across all four conditions, with no significant differences between G-Rex and LimONE-expanded cells regardless of transduction status (Supplementary Figure 2C). This indicates that mitochondrial integrity was preserved across culture platforms and independent of transduction. In contrast, SRC, measured by extracellular flux analysis, was substantially higher in untransduced LimONE cells compared with untransduced G-Rex cells (P < 0.0001), transduced LimONE cells (P < 0.001), and transduced G-Rex cells (P < 0.001) (Figure 5D). Following transduction, SRC in LimONE-expanded cells declined to levels statistically indistinguishable from G-Rex. Despite this reduction in SRC, real-time OCR profiling throughout the Mito Stress Test (basal respiration, oligomycin, FCCP, and rotenone/antimycin A) showed that CAR-T cells transduced in LimONE maintained consistently higher OCR than transduced G-Rex cells at every stage of the assay, including basal and FCCP-stimulated maximal respiration (Figure 5C). Together, these results demonstrate that CAR-T cells manufactured on the LimONE display a favorable phenotype, potent killing activity, and substantially enhanced metabolic fitness, hallmarks of a high-quality CAR-T product.

**Figure 5.**
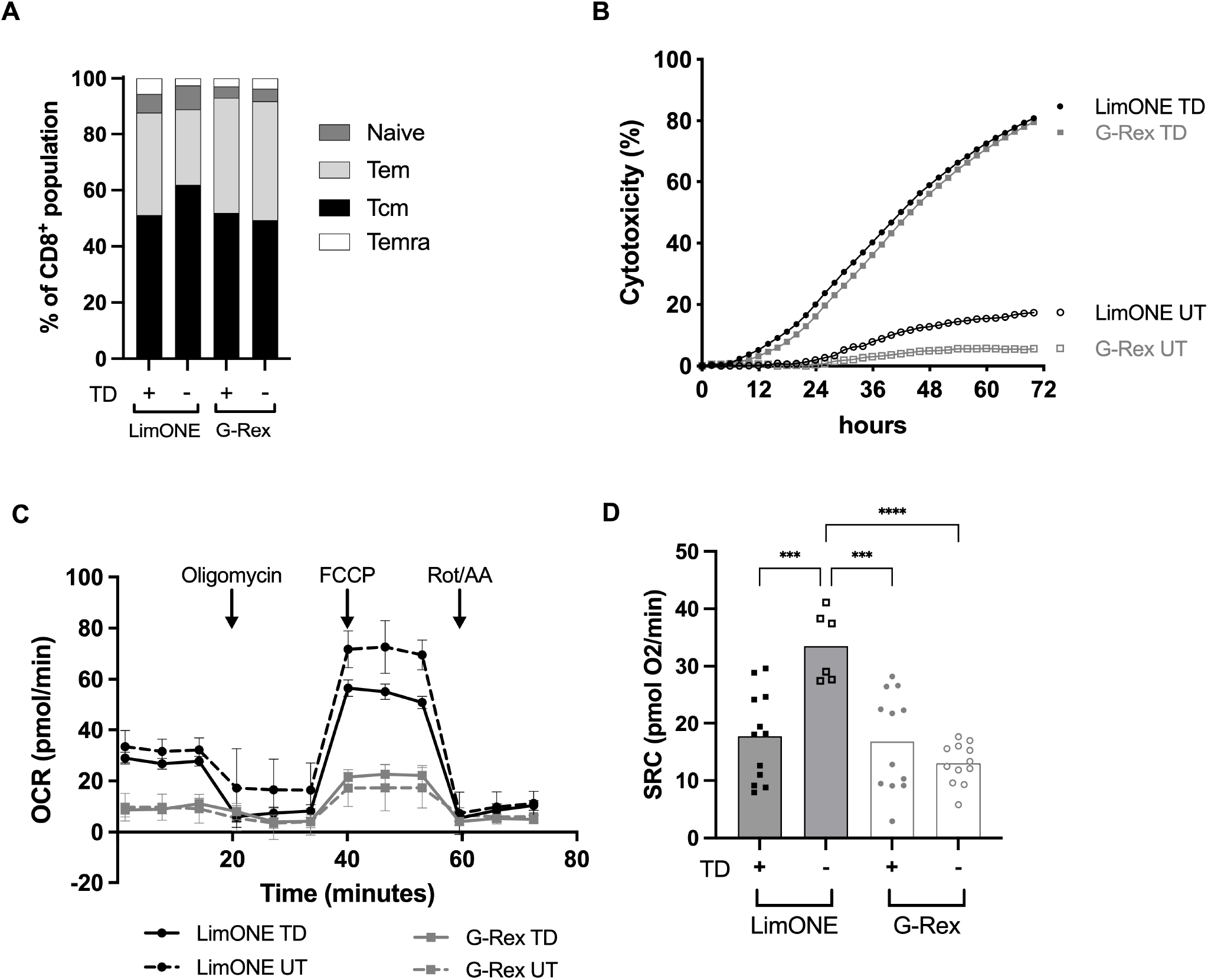
Phenotype, killing activity, and metabolic fitness of LimONE-manufactured CAR-T cells. At harvest, cells were characterized for (A) CD8⁺ differentiation phenotype (Tcm, Tem, naive, Temra) by flow cytometry. (B) Cytotoxicity was assessed over time in triplicate at a CAR-T normalized 1:1 ratio with target NALM6 cells. Finally metabolic activity was determined by measuring (C) Oxygen consumption rate (OCR), with sequential injection of oligomycin, FCCP, and rotenone/antimycin A (Rot/AA) and (D) Spare respiratory capacity (SRC) quantified from (C). Each symbol represents an independent donor, n=4 with triplicates for each donor in panel D. Statistics: (D) ordinary one-way ANOVA with Tukey’s multiple comparisons test. ***p < 0.001, ****p < 0.0001. TD, transduced; UT, untransduced.

### Operational performance and liquid-handling accuracy

Across all four complete CAR-T manufacturing runs, no software or hardware failures occurred. A single process deviation was recorded, caused by an operator who failed to unclamp a line before proceeding with the process. The LimONE software incorporates built-in guardrails to prevent operation outside the instrument’s physical and operational limits. These quality checks included integrity checks on disposables, overflow protection for the LimCORE, backpressure safeguards to protect air filters, and parameter validation to reject out-of-range settings (e.g., incubation duration, centrifugation speed).

The accuracy of automated liquid transfers and of centrifugation-based volume concentration on the LimONE was evaluated (see Methods), with all measurements falling within the ±5% acceptance threshold (Figure 6). For liquid transfers, relative error remained within this threshold for both the 50 mL and 100 mL target volumes tested (n=6 per condition; Figure 6A). For centrifugation-based volume concentration, relative error also remained within ±5% for both starting volumes tested, 350 mL (n=6) and 700 mL (n=4), with end-point volumes closely matching the 50 mL target in both conditions (Figure 6B).

**Figure 6.**
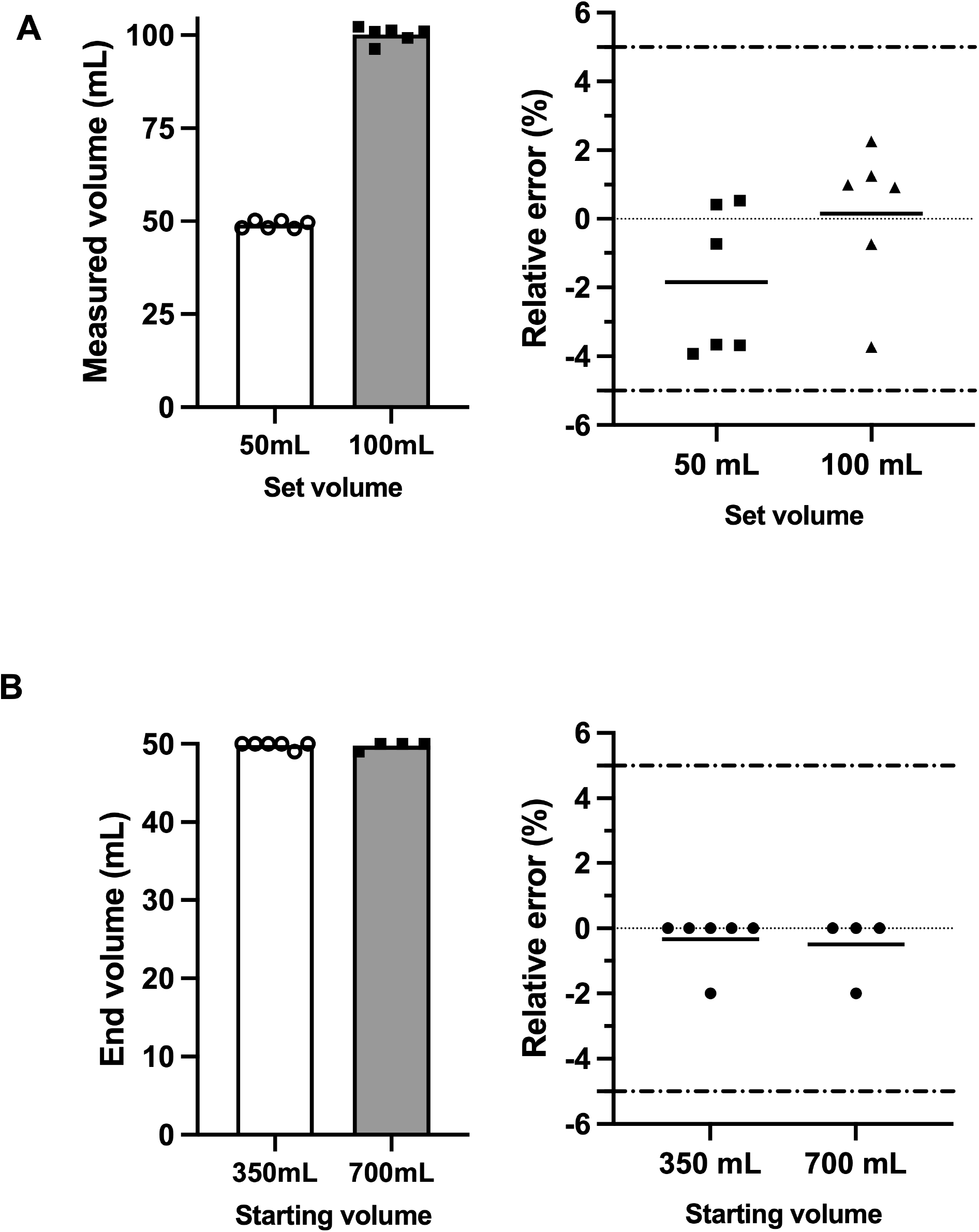
Assessment of liquid transfer accuracy. Accuracy of liquid transfer on LimONE was assessed by testing (A) a target volume (50mL or 100mL) of liquid transfer from a bag to LimCORE. Set volumes were transferred and the volume retrieved in the LimCORE measured (left panel). Relative error (%) of liquid transfers was determined (right panel) with a target of an error <5% (dotted lines). (B) The accuracy of volume reduction via centrifugation was tested. Volumes (left panel) as well as relative error (%) with a target of an error <5% (dotted lines) (right panel) are reported.

## Discussion

Although CAR-T therapies have improved outcomes for many neoplastic and autoimmune diseases, cell therapy predates CAR-T by many decades. Safe and routine blood transfusion was pioneered in the early 20^th^ century after Landsteiner’s discovery of the ABO blood groups, and hematopoietic stem cell transplantation was standardized for clinical practice by Thomas in the 1970s [16,17]. Both therapeutic approaches are now commonly available in hospitals across the world. Importantly, production and control of these therapies have remained in the hands of blood banks and stem cell labs across the world, under a variety of different regulatory structures. Thus, local hospital-based production of complex blood-derived therapies has historical and regulatory precedent. The most recent approval of a hospital-exemption based approach to CD19 CAR-T in Spain is an extension of this approach to local hospital-based production of cellular therapies [18].

Despite the clinical success of CAR-T and other effector T cell therapies, local production has been limited by the complexity of the manufacturing process and the lack of accessible manufacturing solutions. Here we present LimCORE and its implementation from R&D (LimGROW) to automated system designed for GMP manufacturing (LimONE), designed to enable hospital-based, point-of-care manufacturing of cell therapies. Within LimONE, LimCORE serves as the culture vessel, integrated with centrifugation and an automated control system designed to simplify local CAR-T manufacturing without highly trained operators. We show data on the performance of LimGROW and LimONE for T cell selection, expansion, transduction, and precise volume control, with results comparable to manual manufacturing methods using the industry-standard G- Rex bioreactor. Although demonstrated here using a conventional CD19 CAR lentiviral product, the workflow is compatible with increasingly complex engineered cell products incorporating multiplex gene editing, cytokine armoring or TCR engineering.

The current LimONE workflow covers the core manufacturing steps, from cell selection through harvest. Sterile connection of starting material and pre-aliquoted reagents is performed manually. Formulation, fill and finish are planned as the next stage of automation. Formal aseptic process simulation and validation of the fully closed status of the consumable set are planned to support full GMP compliance for use outside highly classified environments. Future studies should evaluate the platform with patient leukapheresis products, clinical-scale manufacturing workflows, and a broader range of engineered cell therapy products, including more complex genetically modified cells.

In*-vivo* gene delivery approaches represent an exciting emerging direction for immune cell engineering, *ex-vivo* manufacturing is expected to remain essential for many applications requiring defined cellular composition, extensive quality control, complex genetic engineering, or precise release testing. These include gene-edited cell products, regulatory T cells, engineered NK cells, TCR-T cells and armored CAR-T products. Furthermore, not all patients and not all diseases are good candidates for in-vivo therapies. For example, in-vivo CAR T therapy in an ALL patients would risk introducing a CAR into a leukemia cell, potentially shielding the target epitope from CAR killing [19]. Other therapies such as Treg-CAR therapies are also poor candidates for in-vivo therapy due to the extreme difficulty of targeting the Treg population and the potential risk of off-target transduction into non-Treg [20]. Large-scale manufacturing of *in-vivo* vectors remains to be demonstrated. Stability issues for viral-like particles or lipid-based particles remain unresolved. Closed *ex-vivo* manufacturing also allows analytic assays to be performed on the cellular drug product, which cannot currently be done with in vivo approaches. Therefore, *ex-vivo* production of cell therapies is likely to continue to be a common therapeutic approach for many years to come.

In conclusion, as production of autologous effector T cell therapies is decentralized and democratized, the availability of reliable and easy-to-use all-in-one manufacturing platforms will be increasingly important.

## Author contributions

Conception and design of the study: CB, YP, JHE, DM. Acquisition of data: CB, LD, FJ, LB, EN, NV, MM, VW, MP, JM, RV, DM. Analysis and interpretation of data: CB, LD, EN, YP, NV, MM, VW, MP, MI, JM, RV, DM. Drafting or revising the manuscript: CB, LD, FJ, LB, EN, YP, JHE, NV, MM, VW, MP, MI, JM, RV, DM. All authors have approved the final article.

## Acknowledgements

The authors would like to acknowledge the Limula engineering team for the design, construction, and ongoing technical support of the LimGROW and LimONE hardware, software, and consumable kits used in this study. The authors would also like to thank Luc Henry for his contribution to funding acquisition and to the supervision of the work. Finally, the authors also thank Akron Biotechnology and Akadeum Life Sciences for providing reagents and for valuable scientific discussions.

## Declaration of generative AI and AI-assisted technologies in the manuscript preparation process

During the preparation of this work, the authors used Claude (Anthropic) for language editing and improving the clarity of the manuscript text. The authors reviewed and edited the output as needed and take full responsibility for the content of the published article.

## Abbreviations

ANOVA: Analysis of variance
AO/PI: Acridine orange/propidium iodide
ATCC: American Type Culture Collection
BACS: Buoyancy-activated cell sorting
BSC: Biological safety cabinet
CAR-T: Chimeric antigen receptor T (cell/therapy)
CD: Cluster of differentiation
FC: Flow cytometry
FCCP: Carbonyl cyanide-4-(trifluoromethoxy)phenylhydrazone
GFP: Green fluorescent protein
GMP: Good manufacturing practice
IL-2: Interleukin-2
MB: Microbubbles
MFI: Mean fluorescence intensity
MOI: Multiplicity of infection
NK: Natural killer (cells)
OCR: Oxygen consumption rate
PBMC: Peripheral blood mononuclear cell
PEI: Polyethylenimine
R&D: Research and development
SD: Standard deviation
SRC: Spare respiratory capacity
Tcm: Central memory T cells
TD: Transduced
Tem: Effector memory T cells
Temra: Terminally differentiated effector memory T cells re-expressing CD45RA
TMRM: Tetramethylrhodamine methyl ester
Tnaive: Naive T cells
UNIFR: University of Fribourg
UNIL: University of Lausanne
UT: Untransduced

---

**Supp Figure 1.**
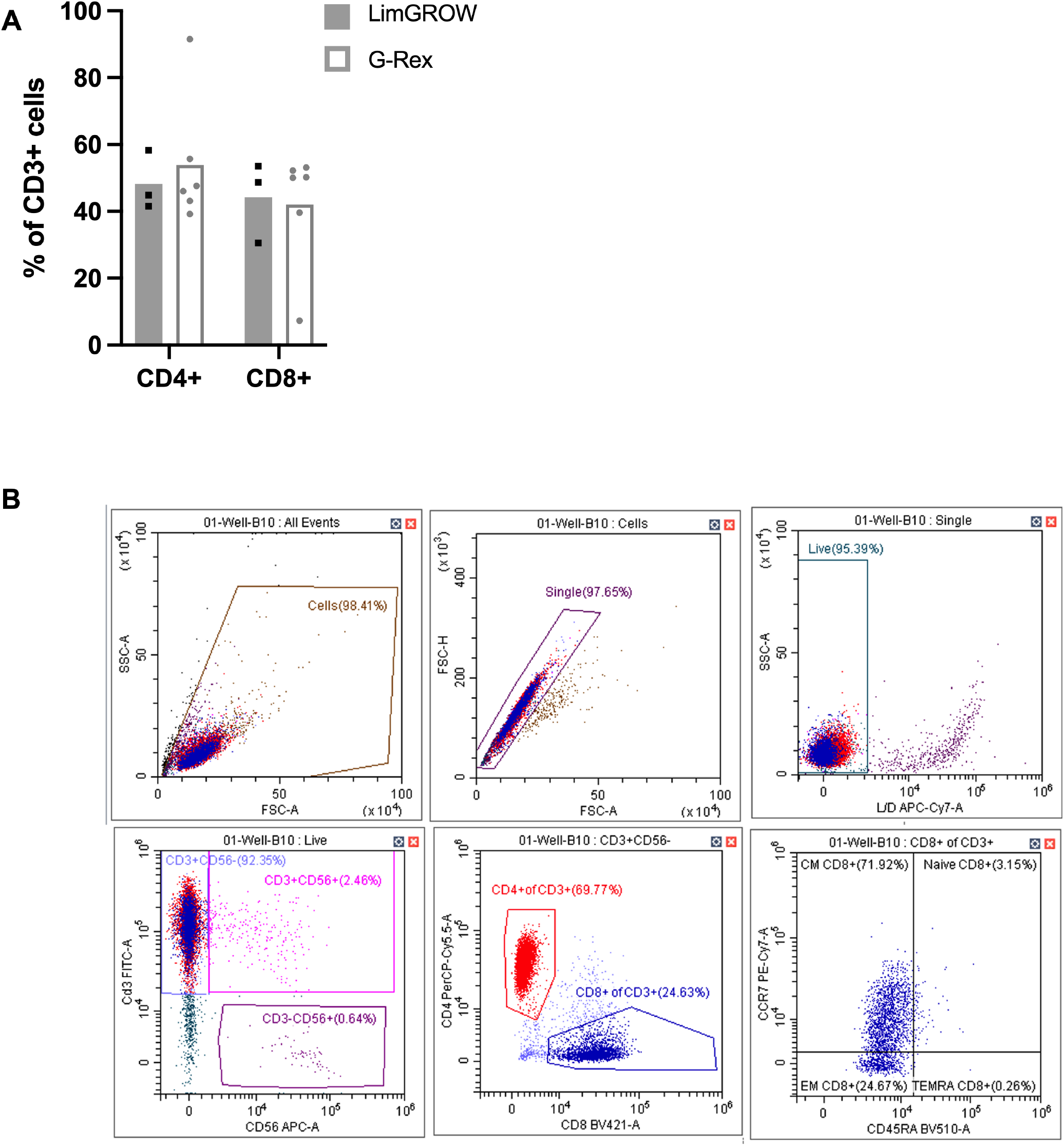
T cell composition and gating strategy. (A) CD4⁺ and CD8⁺ proportion of CD3⁺ cells in LimGROW vs. G-Rex. Each symbol represents an independent donor (n=3 to 6); bars represent mean. (B) Representative flow cytometry gating strategy. Statistics: two-way ANOVA with Tukey’s multiple comparisons test. *p < 0.05, **p < 0.01, ***p < 0.001, ****p < 0.0001; ns, not significant (not displayed).

**Supp Figure 2.**
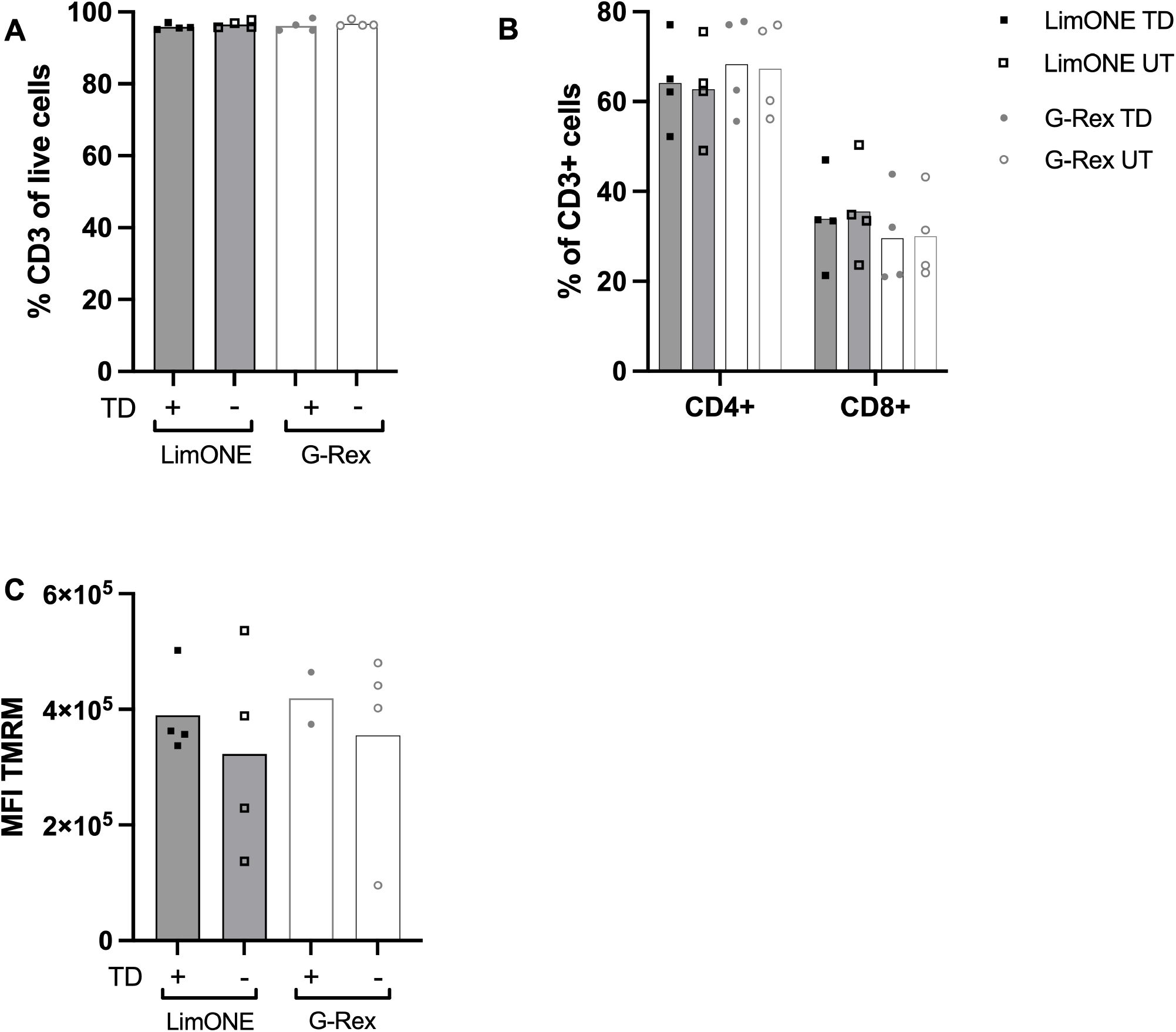
Viability, T cell composition, and mitochondrial membrane potential of LimONE-manufactured CAR-T cells. At harvest, (A) cells viability was determined based on automated AO/PI cell count. (B) CD4⁺ and CD8⁺ proportion of CD3⁺ cells was assessed by flow cytometry as well as (C) Mitochondrial membrane potential (MFI TMRM). Each symbol represents an independent donor (n=4). Statistics: two-way ANOVA with Tukey’s multiple comparisons test. *p < 0.05, **p < 0.01, ***p < 0.001, ****p < 0.0001; ns, not significant (not displayed).

